# The Impact of High-Performance Computing Best Practice Applied to Next-Generation Sequencing Workflows

**DOI:** 10.1101/017665

**Authors:** Pierre Carrier, Bill Long, Richard Walsh, Jef Dawson, Carlos P. Sosa, Brian Haas, Timothy Tickle, Thomas William

## Abstract

High Performance Computing (HPC) Best Practice offers opportunities to implement lessons learned in areas such as computational chemistry and physics in genomics workflows, specifically Next-Generation Sequencing (NGS) workflows. In this study we will briefly describe how distributed-memory parallelism can be an important enhancement to the performance and resource utilization of NGS workflows. We will illustrate this point by showing results on the parallelization of the Inchworm module of the Trinity RNA-Seq pipeline for *de novo* transcriptome assembly. We show that these types of applications can scale to thousands of cores. Time scaling as well as memory scaling will be discussed at length using two RNA-Seq datasets, targeting the *Mus musculus* (mouse) and the *Axolotl* (Mexican salamander). Details about the efficient MPI communication and the impact on performance will also be shown. We hope to demonstrate that this type of parallelization approach can be extended to most types of bioinformatics workflows, with substantial benefits. The efficient, distributed-memory parallel implementation eliminates memory bottlenecks and dramatically accelerates NGS analysis. We further include a summary of programming paradigms available to the bioinformatics community, such as C++/MPI.

## INTRODUCTION

In a recent article Lockwood reported: “There is no reason a high-performance framework for operating a distributed set of DNA sequence reads cannot be similarly developed” [1]. In this article the development of scientific applications in the genomics community was being compared to practices in computational chemistry. Computational chemistry developers tend to make HPC best practices a priority when developing software. Today’s genomics software ecosystem consists of a large number of open source programs that are often developed and adapted to desktop computers, laptops, or even tablets and designed to operate within computing resources confined to the device [1]. Some specialized bioinformatics software packages are developed to utilize a distributed computing and distributed-memory system, with examples including Abyss [2], Novocraft [3], mpiBLAST/Abokia-BLAST [4]-[6] or HMMER 2.3.2 [7], PhyloBayes [8], Ray [9], Meraculous [10] and now Trinity [11]. The continued adoption of HPC best practices by the bioinformatics community will prove to put method development on equal footing with communities such as chemistry, engineering, and physics. These communities have successfully leveraged well-established standards such as Message Passing Interface (MPI) [12], utilized on distributed-memory parallel platforms.

Fundamentally, parallel programming enables an application to utilize more resources, including processor cores, memory capacity, memory bandwidth, and network interconnect bandwidth. This is extremely important, because many important computational problems are simply not tractable when limited to the resources on a single core or a single node. Two commonly used parallel programming modalities are OpenMP and MPI.

OpenMP [13] is a shared-memory parallel programming model. It provides syntax, primarily compiler directives, defining *threads,* which can execute independently and simultaneously. It is typical, for example, for loops to be executed in parallel by allowing each iteration loop to be executed on a distinct thread, and for each thread to be assigned to a single core. All of the threads executing in an OpenMP program have access to all of memory allocated to the job, but no access to memory on other compute nodes within a system. In general, an OpenMP program can use all of the resources (cores, memory capacity, memory bandwidth, interconnect bandwidth) on a single node, but typically cannot use any resources that exist on other nodes.

MPI is the most commonly used distributed-memory parallel programming model and provides a set of functions that enable communication between nodes. In an MPI program, one specifies multiple *ranks*, which by definition execute independently and simultaneously, but can exist on distinct nodes. Most commonly, data is decomposed into independent sections, and processed independently, with relatively infrequent communication between the ranks. Because MPI allows for communication between ranks that reside on different nodes, MPI applications can utilize all of the resources on an arbitrary number of nodes.

Figure 1 illustrates the level of hardware resource availability for shared-memory parallel applications (typically OpenMP), distributed-memory applications (typically MPI), and hybrid parallel applications, which use both shared- and distributed-memory parallelism. Figure 2 shows the association of processors (typically cores) and nodes’ memory for serial, shared-memory parallel, and distributed-memory parallel applications.

**Figure 1.**
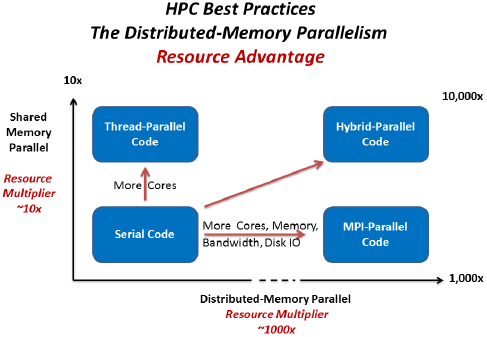
Comparison between distributed- and shared-memory parallelism. Notice the large difference of “resource multiplier “ in the two axis.

**Figure 2.**
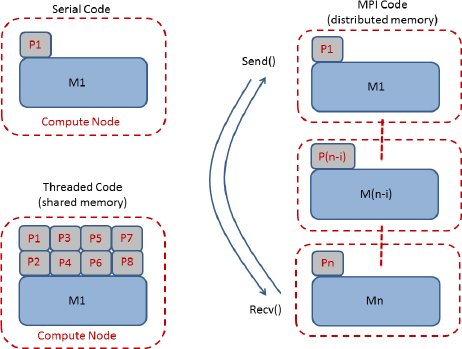
Comparison between computer code implementation, depending on architectures. Notice that MPI and Threaded codes can be combined to form so-called “hybrid MPI” codes, where threads are shared on each node only.

It is important that the HPC community understands the close interplay between the genomics software ecosystem and sequencing technologies. Typical NGS analysis is performed by a selected set of independent programs that need to be joined in a particular order (often with the help of scripts, or workflow managers such as *The Galaxy Project* [14], *Gene Pattern* [15], or *Illumina BaseSpace* [16]) in order to form a specific genomic workflow. *Genomic workflows are the backbone of NGS secondary analysis.* They are flexible and are meant to adapt swiftly to advances in sequencing technologies (such as the *454 Life Sciences* [17], *Illumina* [18], *Pacific Biosciences* [19], or the newer *Oxford nanopore* [20] sequencers). Two examples of genomic workflows are the so-called MegaSeq [21] and Churchill [22] workflows; both calling sequence variants from fastq sequence files by uniquely combining component software including: BWA [23], Picard [24], Samtools [25], and GATK [26]. It is important to realize that each of these component programs performs a specific task that is of limited value on its own; it is however the *combination* of tasks throughout a given workflow that generates the desired solution. Within typical workflows any two intermediary tasks are typically connected through input/output (I/O). This implies a larger amount of read/write I/O in comparison to other more conventional HPC programs in computational chemistry or physics. Nevertheless, bioinformatics I/O formats, starting from the sequencer’s output to the resulting “variant calling”, is relatively standardized and, if reduced to a minimum, may not necessarily represent a significant bottleneck to a HPC implementation.

Finally, to understand how the NGS community can embrace HPC best practice, it is important to describe the current factors that differentiate these two communities, namely computational chemistry and NGS. As previously pointed out [1], method implementation in computational chemistry has evolved together with HPC. In the late 1980’s, the performance gap between “departmental” machines and state of the art HPC systems was at least a factor 100X. This provided a very strong incentive to move implementations to a HPC system. Conversely, NGS software development has evolved by reacting to the prevailing sequencing technologies. This technology is rapidly and continually changing and forces developers to focus on functionality rather than system size [1].

In order to embrace HPC best practice in such a dynamic field as NGS, it is important for HPC software development to evolve within the ecosystem. Upon embracing HPC, the same HPC best practices used in computational chemistry can be applied to entire NGS workflows, including workload distribution, compute parallelization, and system profiling to optimize resource utilization and eliminate bottlenecks.

As an example of how HPC best practices can be applied in an NGS application, here we describe our efforts to integrate runtime and memory scaling into the Inchworm module of the Trinity *de novo* assembly software. We introduce minimal changes to the sequential version of Inchworm, yielding the first generation of a distributed-memory parallel Inchworm (termed MPI-Inchworm). MPI-Inchworm is demonstrated to leverage distributed-memory systems and to scale computing across hundreds to thousands of compute cores. We expect there should exist ample opportunities to apply HPC best practices in other NGS workflows, leveraging techniques described here in our development of MPI-inchworm.

This paper is divided as follows: Section I contains a discussion of the main features of HPC best practices, applied to bioinformatics; Section II describes the original shared-memory single-node version of Trinity; Section III is the core of this article where we focus on the details of the distributed-memory multi-node implementation of Inchworm and how it relates to HPC best practices; Section IV describes the hardware leveraged here; Section V reports results associated with the two concepts of time-scaling and memory-scaling, as defined in Section I on HPC best practices; and Section VI details concluding remarks.

## I. HPC BEST PRACTICE

A core concept underlying HPC best practices is distributed-memory parallelism (described in Figure 1), which implies two important HPC features: time scaling and memory scaling. Time scaling helps reduce the time-to-solution and is essential in clinical genomic settings in which medical practitioners are in need of fast and accurate diagnostics. Simply using embarrassingly parallel programming workflows and leveraging supercomputers was shown capable of aligning reads and calling variants for 240 human genomes in just over 50 hours [21]. Another application is demonstrated capable of similar alignment and variant calling in less than two hours per sample [22]. Time-scaling remains indeed one of the most fundamental features of HPC, and potentially most relevant in clinical applications of NGS analysis.

The second HPC focus, memory scaling, is by no means less important, but often neglected. Memory constraints are often an issue on shared-memory nodes (e.g., a typical desktop), because large NGS problems can easily require more memory than is available on a typical single node. Excellent parallel threading (i.e., time scaling) can be achieved on a single node using, for example, OpenMP [13]. Unfortunately, such parallel threading can never solve the problem of being restricted to a single shared memory. On distributed-memory architectures, however, once the data for assembling or aligning a DNA/RNA sequence is distributed among typical (low-cost) microprocessors, the memory limitation no longer imposes an inherent constraint. Such memory scaling, in principle, enables HPC computation on any genome size, from the modest human genome at ∼3.2 Gb (giga base pairs or billion base pairs) to the spruce (∼20 Gb), or even larger genomes that have not yet been sequenced, such as the *axolotl* (∼30Gb), or *Paris japonica* (∼149 Gb) - the Japanese flower estimated to have the largest estimated genome size [27]. Moreover, the ability of a single program to use the memory of multiple nodes allows systems to be configured homogeneously, rather than requiring a number of expensive large-memory nodes for special cases.

In practice, distributing the data might require refactoring certain parts of an existing serial implementation. Although the concept of message passing (MPI) is straightforward, it does involve added computational complexity and requires developers to overcome certain barriers. This can easily be achieved by becoming familiar with MPI [12]. In the case of Trinity, the challenges in integrating MPI for distributed-memory allocation and distributed computational processing involved deciding exactly how to distribute the input sequencing reads and deciding which data needed to be communicated among the distributed computations. Our effort to integrate MPI into the Inchworm component of the Trinity software is further discussed in Section IV.

What exactly are HPC best practices? Figure 3 shows a diagram depicting steps to guide one in applying HPC best practices. Evaluations start with I/O; any unnecessary I/O occurring inside a particular task of the workflow should be eliminated, keeping only the I/O that is necessary (i.e., keeping essentially only the I/O situated at the beginning and the end of a given task inside that workflow). The second item, memory usage, is related to the I/O; ideally input data files are read in parallel by each MPI rank and then data is distributed among the ranks to best support the program algorithm. Once distribution of the data is optimized, the CPU usage and communications can be evaluated to further improve time-scalability. The last two items, CPU time and communication patterns, are not generically approached but often require solutions more specific to the algorithm itself. We summarize issues relating to HPC best practices below: (comparing with Figure 3).

**Figure 3.**
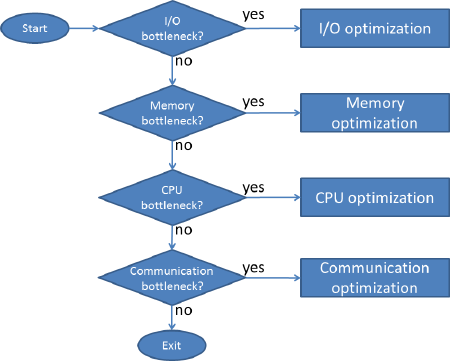
HPC best practices diagram

Profiling is essential to HPC best practices. This process consists of timing each important step of any application; separating the time spent on the I/O from the time spent computing, locating code constructs that are slower than others, and determining how much memory is used. Profiling should always be included in any complex application. There are two major ways to implement profiling. First, including the logic to perform profiling directly within your code controlled via command-line options or compiler flags (e.g., a “PROFILE” logical flag that can be turned on and off in the Makefile), with the help of libraries such as <ctime> (time.h), or MPI_wtime() for timing sections of code, or alternatively, using ‘third party’ software tools such as Cray PerfTools [28], Tau [29], or Collectl [30]. Once the serial application is thoroughly profiled, then the developer can begin parallelizing the application, such as by leveraging MPI.

Below, we show an example of a small MPI program written in C, which demonstrates two key features of an MPI program: (1) each rank can execute specific sections of code as determined by its rank assignment, and (2) different ranks can communicate by sending/receiving messages. This program runs as two distributed parallel processes where process rank 1 uses MPI communication to send a message containing the nucleotide sequence ‘GATTACA’ to process rank 0, which simply reports the message it received.

Incorporating MPI into any parallel C/C++ program requires a minimum of six extra lines, as shown below in our example program’s “SendDNA.c” file, see [31]-[33] and some extra useful examples of parallel sorting in [34]:

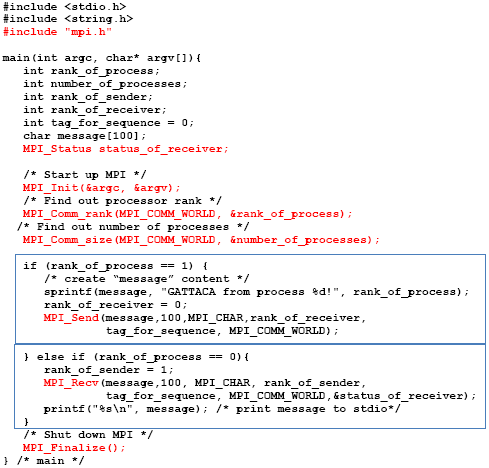

This program is compiled using:

~~~
CC -o ./SendDNA SendDNA.c
~~~

A batch job contains the following line:

~~~
mpirun -n2 ./SendDNA
~~~

where “-n 2” specifies the number of MPI ranks. The result is:

~~~
GATTACA from process 1!
~~~

which is actually output by the MPI process with rank 0.

This minimalistic MPI program shows the six required lines, marked in red, that are exclusively related to MPI (in addition to the code inside the two boxes that define the communication pattern). Each of the two ranks is running as a separate process and is aware of its individually assigned rank number. Based on its rank, each process will need to know what its role is and what data it should be responsible for processing and which sections of code to execute. The different ranks communicate messages with each other via MPI_Send() and MPI_Recv(), as shown schematically in Figure 2, and as implemented in the program above. In our example, the code in the top box is only executed by rank 1, putting the message “GATTACA from process 1” into the variable *message,* and then sending this message to rank 0. The code in the bottom box is only executed by rank 0 residing at a different compute node, where it receives the message from rank 1, stores it in its local variable *message*, and prints the message to standard output.

Finding the best approach for distributing data among MPI ranks is of fundamental importance, because it will also determine the amount of communication incurred by the algorithm. Having the data properly distributed allows the total memory footprint at each compute node to diminish with increasing number of nodes or MPI ranks. The memory footprint can also be profiled and will be discussed in section IV. Once proper scalable memory distribution is achieved then one can work on parallelizing the algorithm for computing on these data. This requires practical testing as much as theoretical algorithmic evaluation. Throughout this process of developing a parallel application it is imperative to keep in mind the diagram of HPC best practices in Figure 3.

The next two sections show how we applied HPC best practices to the parallelization of Trinity-Inchworm.

## II. TRINITY RNA-SEQ

Trinity [11] is a popular *de novo* RNA-Seq assembly tool that reconstructs transcript sequences from RNA-Seq data and, unlike other related popular methods such as the Tuxedo tool suite [35], Trinity does not require a reference genome sequence. Trinity, as the name suggests, consists of three main components: Inchworm, Chrysalis, and Butterfly. These three components form a workflow that is united by a Perl script called Trinity.

Briefly, the components of Trinity operate as follows. Inchworm builds a catalog of k-mers (sequence of bases of length k, default of k=25) from all the reads such that every k-mer is associated with its frequency of occurrence within the full set of reads. The single ‘seed’ k-mer with the greatest occurrence is selected from the catalog, and a contig is constructed by greedily extending the seed from each end, selecting overlapping k-mer that has the highest occurrence and extends the growing contig by a single base. Single base extensions continue to occur until no k-mer exists in the catalog that can yield an extension, in which case the resulting contig sequence is reported by Inchworm, and the k-mers comprising the reported contig are eliminated from the k-mer catalog. Rounds of seed selection and contig extension continue until the k-mer catalog is exhausted.

The remaining steps of Trinity involve Chrysalis clustering Inchworm contigs that are related due to alternative splicing [Inchworm contigs sharing (k-1)-mers], and building a de Bruijn graph for each cluster, with ideally one cluster of contigs per gene and a corresponding de Bruijn graph representing the transcriptional complexity exhibited by that gene. The Butterfly software then threads the original reads through the de Bruijn graphs and reconstructs the full-length transcript sequences and splicing patterns best represented by the reads in the context of the graph.

Trinity was initially designed as a single node large memory application, where only the final phase involving Butterfly was embarrassingly parallel and could be computed using a distributed computing environment. Profiling of the original Trinity code using Collectl was performed and published previously in [36]. Those profiles have shown improvements in terms of shared-memory parallelism, using up to 32 CPU cores. Most importantly, profiling has identified time and memory bottlenecks located in the Trinity workflow. Those bottlenecks were shown to exist at specific sections of the Inchworm and Chrysalis codes.

Here, we engineered a version of the Inchworm software to scale efficiently, both in terms of memory footprint and wall clock time, leveraging MPI. (MPI-based parallelization of Trinity’s Chrysalis module has also been implemented by a separate group as a separate effort [37].) The next section describes our implementation of the distributed and massively parallel version of Inchworm we named MPI-Inchworm.

## III. DEVELOPMENT OF MPI-INCHWORM

The general structure of MPI-inchworm contains two main phases, a k-mer Distribution Phase (Fig 4a) and a Contig Building Phase (Figure 4b). The first phase populates the k-mer catalog in distributed-memory. The second phase builds the contigs via the seeded k-mer greedy extension algorithm described earlier. The number of ranks can easily vary from 1 (not distributed) to thousands of ranks (distributed). The input data and the assembly computations are equally distributed among all of the ranks. Below, we describe the overall architecture of MPI-Inchworm and how we distribute the data and computations.

**Figure 4.**
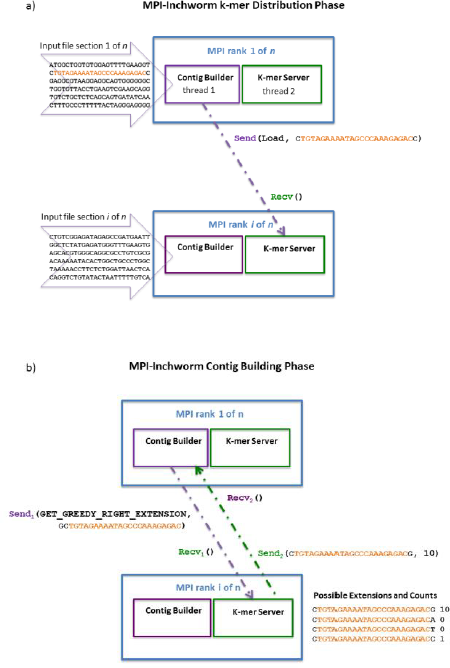
Parallel Inchworm algorithm (a) Phase 1: K-mer distribution (b) Phase 2: Contig building

Each distributed MPI rank is composed of two internally parallel OpenMP threads that further subdivide the work at each MPI rank. In Figure 4, each distributed MPI rank is depicted as a box with a blue outline, and the two internally parallel threads are shown as the ‘contig builder *(cb)’* thread (purple outline) and ‘k-mer server *(ks)’* thread (green outline). In describing the distributed computations, we make reference to the role of each of these threads within a rank. In general, the *cb* thread of one rank sends a message to the *ks* thread of another rank, and that *ks* thread then responds accordingly.

In the initial k-mer Distribution Phase (Figure 4a), MPI-Inchworm parses the input RNA-Seq reads, extracts the k-mers from the read sequences, and stores each k-mer in distributed memory. Here, MPI-inchworm employs the parallel I/O programming model. The input file is subdivided into as many sections as there are MPI ranks requested for a particular run, and each MPI rank reads and processes only its assigned section of that file. The *cb* thread is responsible for parsing the reads, extracting the k-mers, and defining the location where the k-mer will be distributed. In addition to each rank reading a section of the reads file, each rank is responsible for locally storing k-mers, however, the k-mers being extracted by a given rank are not necessarily stored locally by that rank. Each k-mer inherently encodes the identity of the rank that is responsible for storing it by virtue of its nucleotide composition. By extracting the central sequence of a k-mer (excluding the base at each terminus), the destination rank is identified. This simply involves converting the k-mer sequence to an integer value, and dividing it by the total number of ranks, with the value of the remainder used to identify the rank at which the k-mer is to be stored. If the destination rank is the same as the rank to which the *cb* thread is assigned, then the *cb* thread stores that k-mer locally. Otherwise, the *cb* thread uses MPI to send (MPI_Send) the k-mer to the destination rank, where the corresponding destination *ks* thread will receive (MPI_Recv) the k-mer and store it at that location. In addition to storing the k-mer, the frequency of that k-mer is stored. If the k-mer already exists at a given location, then its count is incremented to reflect the additional occurrence. This k-mer distribution pattern ensures that the memory footprint decreases with an increase number of nodes, as discussed in Section II on HPC best practices.

Once all the k-mers have been stored in distributed memory, the subsequent Contig Building Phase can begin. In this phase, each rank builds a different contig according to the Inchworm greedy extension algorithm, with slight modifications^1^. Starting with a seed k-mer, the cb thread will search for a greedy extension. Any of the four possible singlebase extensions to the seed (G, A, T, or C) would involve an extension k-mer containing the same central sequence, and given that the destination rank for a k-mer is based on the central sequence (described above), all four possible extension k-mers would be co-located at a single destination rank. If the destination rank is that of the *cb* thread, then the *cb* thread will look up the greedy extension k-mer locally. Otherwise, the *cb* thread sends a message to the rank containing the extension k-mers, requesting the greedy extension. The *ks* thread at the destination rank receives the request, identifies the four possible extension k-mers and responds with that k-mer that is most frequent, or with a message that no such k-mer was available. Once the *cb* thread completes building a contig, it then sends ‘delete k-mer’ messages to destination ranks so that those k-mers are removed from the fully distributed k-mer catalog.

This process of seed selection and contig extension is repeated throughout all the MPI ranks. As each rank constructs contigs, it writes the contig sequences to a rank-specific output file. After all ranks have exhausted their local k-mer stores, the MPI-Inchworm contig construction phase is complete. A final “harvesting” (serial) routine operates to consolidate the contigs output from each of the ranks, remove any redundancy, and prepare a single output file to be used as input to Chrysalis, the next step of Trinity.

In terms of HPC best practices, communication has been minimized. This was done through examining profiling data, as shown in Figs. 5 and 6. The entire communication pattern in MPI-Inchworm essentially requires two MPI commands: MPI_Send and MPI_Recv, as in our earlier example code. Additional information about the MPI inchworm implementation can be found in the following recorded seminar presented by author BH [38].

**Figure 5.**
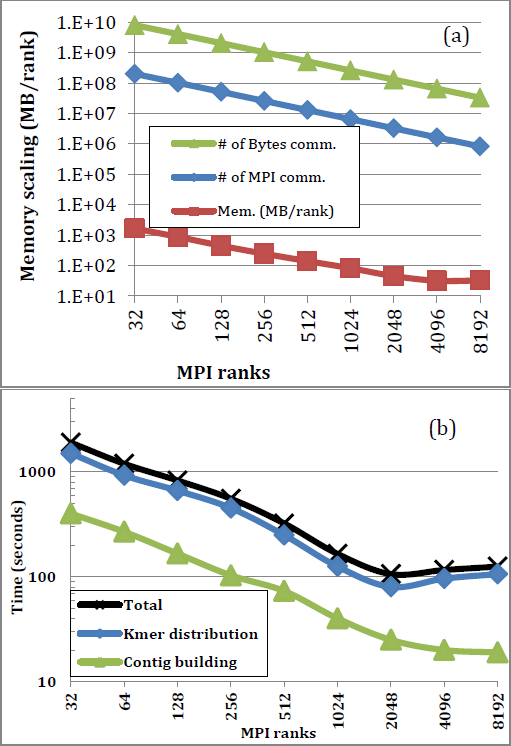
(a) Memory scaling and (b) Time scaling of MPI-Inchworm on the mouse RNA-Seq data. Computations are done on XC40 Haswell-16, 2 sockets (32cores/node; 128GB/node).

**Figure 6.**
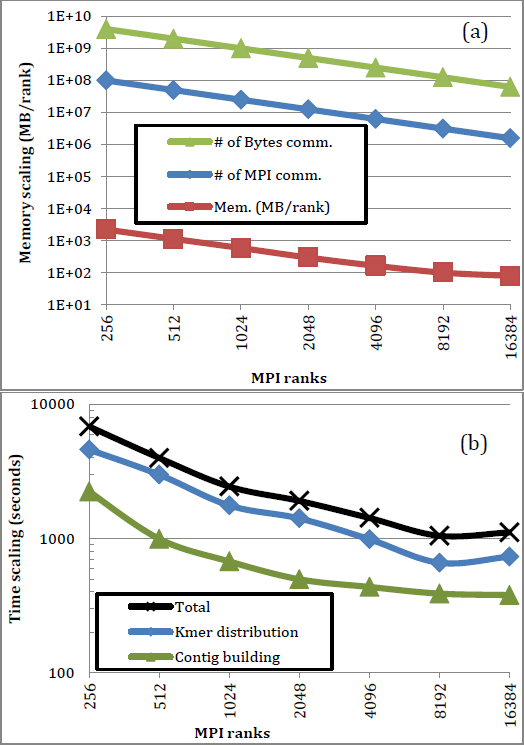
(a) Memory and (b) Time scaling of MPI-Iinchworm on the axolotl RNA-Seq data. Computations are done on XC40 Haswell-16, 2 sockets (32 cores/node;128GB/node).

## IV. HARDWARE RESOURCES

The primary system utilized in this study was a Cray XC40. The processor is a 64-bit Intel® Xeon® E5-2698 V3 “Haswell” 16 core 2.3 GHz processor. There are two processors per compute node and 384 processors per cabinet. The processor peak performance per core is 36.8 GF. The memory consists of 128 GB DDR4-2133 MHz per compute node. Memory bandwidth is 120 GB/s per node. The system interconnect is Cray Aries multilevel dragonfly topology, which provides a low latency, high bandwidth network. There is one Aries router ASIC per four compute nodes. Each Aries has 40 external network ports over 3 levels, providing more than 10 GB/sec bidirectional point-to-point bandwidth, with less than 1.5 μs latency.

## V. RESULTS AND DISCUSSION

In this section we discuss time scaling and memory scaling for two relevant transcriptome data sets corresponding to mouse and axolotl (Mexican salamander), both organisms of importance to biomedical research. The human genome and the mouse are similar in size (∼3 Gb), and the mouse is an important model organism used in many contexts including clinical cancer research. The axolotl is known for its extraordinary regenerative abilities in reconstituting limbs, retina, liver, and even minor regions of the brain [39]. The axolotl’s genome contains about 10 times as many base pairs as that of the mouse or the human, and has yet to be sequenced and assembled due to its massive size and the related cost and complexity of such an effort. The RNA-Seq data leveraged for each organism is described below.

Figure 5 and 6 summarize the (a) memory scaling and (b) time scaling results for MPI-Inchworm on the mouse and axolotl RNA-Seq data, respectively. The memory usage clearly scales with the number of MPI ranks (Figures 5a and 6a). The importance of memory scaling cannot be over-emphasized. For example, the input fastq file for the mouse contains approximately 50 million 76 base length paired-end Illumina RNA-Seq reads. In addition to storing k-mers, additional memory is needed at each node to reconstruct contigs. We find that the mouse can easily run on one 128-GB node with two Haswell-16 processors. Figure 5 shows scaling results for the mouse as performed using 1 node (32 MPI ranks/node) up to 256 nodes on the Cray XC40. In contrast to the mouse data, the axolotl data set consists of ∼1.2 billion 100 base length paired-end Illumina RNA-Seq reads, and this very large data set requires a minimum of 8 nodes *just to have adequate memory capacity*. Figure 6 shows scaling results for the axolotl performed on 8 nodes up to 512 nodes. Out-of-memory errors occur if one tries to run the axolotl’s RNA-Seq on a typical single 128-GB XC40 Haswell-16 node. This is one of the most striking, and often under recognized, advantages of distributed-memory parallelism: memory scalability. Most importantly, the original (non-distributed) Inchworm program can only run jobs on single node, and therefore, cannot run the axolotl’s RNA-Seq on commodity hardware. *Distributed-memory parallelism allows researchers to do research that would otherwise not be possible.*

The MPI-Inchworm algorithm demonstrates excellent time scaling properties up to 2048 MPI ranks (64 nodes) with the mouse RNA-Seq (Figure 3b), and up to 8192 MPI ranks (256 nodes) with the axolotl RNA-Seq (Figure 4b). The larger the RNA-Seq data set, the better the time scaling. We are currently exploring how to improve scalability beyond 2048 MPI ranks and 8192 MPI ranks for the mouse and axolotl datasets, respectively. Overall, the memory and time scaling are excellent up to multiple thousands of MPI ranks. Time scaling is of fundamental importance for instance in clinical environments, where time-to-solution could have a critical impact on patients’ recovery.

## VI. CONCLUSIONS

We have shown how HPC best practices can be applied to the parallelization of an important component of the Trinity RNA-Seq application, the Inchworm contig assembler. The distributed MPI-Inchworm application is now up to 18 times faster on 128 nodes (4096 MPI ranks) than on a single node (i.e., using still 32 MPI ranks on that single node) and can handle data sets that are much larger than what the original non-distributed code is capable of processing, as a result of applying HPC best practices to NGS analysis. In general, we believe that any bioinformatics workflow can greatly benefit from HPC, in terms of time-to-solution as well as enabling new research. Distributed-memory parallelism allows researchers to complete research that otherwise would not be possible given the limitations of commodity hardware.

## ACKNOWLEDGMENTS

We thank Cray Inc. for the computing time on the XC30/XC40 marketing machine. We thank Broad Institute principal investigator Aviv Regev for generously supporting Trinity development efforts and related activities. Finally, we thank Jessica Whited at the Brigham Regenerative Medicine Center for access to the axolotl RNA-Seq data used in our performance studies.

Research reported in this publication was supported by the National Cancer Institute of the National Institutes of Health under Award Number 1U24CA180922-01. The content is solely the responsibility of the authors and does not necessarily represent the official views of the National Institutes of Health.

Author contributions are as follows: PC performed software performance analytics and drafted the initial version of the manuscript. BH engineered MPI-Inchworm based on input from all authors, and assisted in writing the final manuscript. All authors contributed to and approved the final version of this manuscript.

Our MPI-Inchworm program is open source and freely available within the current distribution of Trinity [40]. The source code MPIinchworm.cpp (located in the directory “`Inchworm/src`” of the current distribution) needs to be compiled separately using the Makefile template (`make -f MPI_cray.Makefile`). The name of the executable is “`MPiinchworm`” and can be used with parameters equivalent to the original `Inchworm` software.

## Footnotes

[1] Seed selection follows a two-phased approach. First a random k-mer is selected from the k-mer store at the corresponding rank. A greedy extension is performed to identify a k-mer with maximum abundance, and that k-mer is chosen as the proper seed for Inchworm contig extension.

